# Restructuring of the Immune Contexture Improves Checkpoint Blockade Efficacy in Murine Lung Cancer

**DOI:** 10.1101/2020.10.09.332387

**Authors:** Nancy D. Ebelt, Edith Zuniga, Monica Marzagalli, Vic Zamloot, Bruce R. Blazar, Ravi Salgia, Edwin R. Manuel

**Affiliations:** Department of Immuno-Oncology, Beckman Research Institute of the City of Hope, Duarte, CA; Division of Blood and Bone Marrow Transplantation, University of Minnesota Medical School, Minneapolis, MN; Department of Medical Oncology and Therapeutics Research, City of Hope National Medical Center, Duarte, CA

**Keywords:** Non-small cell lung cancer, immune checkpoint blockade, anti-PD-1, anti-CTLA4, *Salmonella typhimurium*, small-hairpin RNA, indoleamine 2,3-dioxygenase

## Abstract

Therapeutic options for non-small cell lung cancer (NSCLC) treatment have changed dramatically in recent years with the advent of novel immunotherapeutic approaches. Among these, immune checkpoint blockade (ICB), using monoclonal antibodies, has shown tremendous promise in a small proportion of patients. In order to better predict patients that will respond to ICB treatment, biomarkers such as tumor-associated CD8^+^ T cell frequency, tumor checkpoint protein status and mutational burden have been utilized, however, with mixed success. In this study, we hypothesized that significantly altering the suppressive tumor immune landscape in NSCLC could potentially improve ICB efficacy. Using sub-therapeutic doses of our *Salmonella typhimurium*-based therapy targeting the suppressive molecule indoleamine 2,3-dioxygenase (shIDO-ST) in tumor-bearing mice, we observed dramatic changes in immune subset phenotypes that included increases in antigen presentation markers, decreased regulatory T cell frequency and overall reduced checkpoint protein expression. Combination shIDO-ST treatment with anti-PD-1/CTLA-4 antibodies enhanced tumor growth control, compared to either treatment alone, which was associated with a significant intratumoral influx of CD8^+^ and CD4^+^ T lymphocytes. These results suggest that the success of ICB therapy may be more accurately predicted by taking into account multiple factors such as potential for antigen presentation and frequency of suppressive immune subsets in addition to markers already being considered. Alternatively, combination treatment with agents such as shIDO-ST could be used to create a more conducive tumor microenvironment for improving response rates to immunotherapy.

## INTRODUCTION

Non-small cell lung cancer (NSCLC) tumors contain large percentages of T cells (25-46% of the CD45^+^ fraction), many expressing tumor antigen-specific T cell receptors (TCRs) [1, 2]. However, most of these T cells lack effector function and are prone to activation-induced apoptosis due to chronic stimulation by tumor-derived antigens [3, 4]. An important marker of these dysfunctional T cells is programmed death-1 (PD-1), a cell surface receptor that can be activated to cause T cell apoptosis by tumor cells or other immune cells expressing programmed death-ligand 1 (PD-L1) [5-7]. Antibody-mediated inhibition of the PD-1/PD-L1 axis is a promising new treatment option for non-small cell lung cancer (NSCLC), which continues to be the leading cause of cancer-related death in the United States and worldwide [8-10]. Several anti-PD-1/PD-L1 antibodies, including pembrolizumab, nivolumab, and atezolizumab, have been approved for the treatment of advanced NSCLC after remarkable improvement in median overall NSCLC patient survival was demonstrated in clinical trials [11, 12].

The presence of another cell surface molecule, cytotoxic T lymphocyte-associated protein 4 (CTLA-4), suppresses T cell responses by acting as a homolog for CD28 that competes for binding to activating MHC co-receptors present on antigen presenting cells (APCs) [13]. Treatment of NSCLC patients with antibodies targeting CTLA-4 (e.g. ipilumumab) in conjunction with chemotherapy results in modest increases in immune-related, progression-free survival, but overall survival is unchanged [14]. Greater benefit was seen with the combination of ipilimumab and nivolumab, which showed significantly improved overall survival compared to chemotherapy and nivolumab alone [15], resulting in recent FDA approval for frontline treatment of metastatic NSCLC with the ipilimumab and nivolumab combination. These cell surface molecules, and a few others, constitute the immune “checkpoint” proteins. Although the magnitude and durability of responses induced by immune checkpoint blockade (ICB) are unrivaled by current treatment options, only a small percentage of cancer patients will experience clinical benefit [16, 17], indicating mechanisms of resistance to checkpoint blockade that must be alleviated before responses can be improved.

Two major cell types have been connected to checkpoint blockade failure: regulatory CD4^+^ T cells (Tregs) and myeloid derived suppressor cells (MDSCs) [18-20]. Tregs and MDSCs contribute to a suppressive microenvironment through distinct and overlapping mechanisms. Tregs, found in tumor tissue and draining lymph nodes, have high expression of CTLA-4 and Lag-3 that facilitate suppression of dendritic cells (DCs) and T cells through direct cell-to-cell contact; they also secrete immunosuppressive cytokines such as IL-10, transforming growth factor β (TGF-β), and IL-35, which downregulate IFNγ responses, inhibit antigen presentation by APCs, and impair T cell proliferation [21]. MDSCs suppress T cell activation and proliferation through upregulation of reactive oxygen species (ROS) and arginase, as well as indoleamine 2,3-dioxygenase (IDO) [22-24] which suppresses T-cell function by converting essential tryptophan into suppressive kynurenines [25]. MDSCs can also suppress NK cytotoxic function [26], inhibit antigen presentation by APCs, and induce formation of Tregs through secretion of IL-10 [27, 28]. Finally, MDSCs express significant levels of PD-L1, PD-1 and CTLA-4 that can also interfere with T cell activation [29].

Recently, a novel subset of anti-tumor myeloid cells have been identified in the early stages of NSCLC that were capable of inducing anti-tumor immunity through attainment of antigen presentation capabilities [30-33]. Our recent studies in pancreatic ductal adenocarcinoma (PDAC) have also shown that depletion of neutrophils and monocytes abrogate the anti-tumor effects of anti-PD-1 treatment, demonstrating the importance of tumor-associated myeloid cells (TAMCs) in checkpoint blockade efficacy [34]. Increasing antigen stimulation of T cells through increased presentation by APCs and/or converted MDSC may sensitize permanently dysfunctional T cells to ICB therapy. Significantly decreasing the frequency of various immunosuppressive subsets may also be necessary to create a tumor microenvironment (TME) more conducive for ICB.

In previous studies employing models of melanoma, colorectal cancer, and PDAC, we confirmed that our *Salmonella typhimurium*-based platform targeting IDO (shIDO-ST) is specifically engulfed by TAMCs and induces their expansion as well as hyperactivation of their anti-tumor functions [22, 35]. In this study, we confirm the ability of sub-therapeutic doses of shIDO-ST to enhance antigen presentation capabilities in TAMCs, decrease the frequency of suppressive immune subsets, and reduce immune checkpoint expression, ultimately improving efficacy of ICB in a pre-clinical model of NSCLC. Our ability to demonstrate synergy between shIDO-ST and ICB treatment in pre-clinical models of NSCLC may support their combination as a strategy to enhance clinical response rates in patients [36, 37].

## MATERIALS AND METHODS

### Animals and cell lines

C57BL/6 mice were obtained from breeding colonies housed at the City of Hope (COH) Animal Research Center, and for all studies, handled according to standard IACUC guidelines (approved IACUC protocol 17128). The Lewis Lung Carcinoma (LLC1) cell line was obtained from ATCC® (CRL-1642). Cells were maintained in RPMI media containing 10% FBS, 2mM L-glutamine and pen/strep. Prior to subcutaneous implantation into C57BL/6 mice, cells were passaged ≤5 times and maintained at ≤80% confluency.

### *Salmonella typhimurium* (ST)

YS1646 was obtained from ATCC® (202165™). YS1646 was cultured in modified LB media containing MgSO4 and CaCl2 in place of NaCl. The pAKlux2 plasmid was a kind gift from Attila Karsi (Department of Basic Sciences, College of Veterinary Medicine, Mississippi State University, Mississippi State, MS; Addgene #14080). ShScr and shIDO plasmids (Sigma)[35] were electroporated into ST strains using a BTX electroporator (1mm gap cuvettes, settings: 1.8kV, 186 ohms), spread onto LB plates containing 100 µg/mL ampicillin and incubated overnight at 37°C.

### ST administration and neutrophil isolation in tumor-free mice

For blood and spleen neutrophil isolations, on the first day mice were administered 5×10^6^ cfu of shScr, shIDO-ST or HBSS volume equivalent followed by 2×10^6^ cfu on the second and 1×10^6^ cfu on the third consecutive day intravenously via the retro-orbital vein. Blood and spleens were collected after euthanization 48 or 72 hours after the third ST administration. For peritoneal neutrophil isolation, 4×10^7^ cfu of shScr, shIDO-ST or HBSS volume equivalent was injected into the intraperitoneal cavity of mice followed by 2×10^7^ cfu on the second day. Three hours after the second administration, mice were sedated and 5mL of HBSS was injected into the intraperitoneal cavity; the abdomen was massaged to dislodge cells into the fluid. Then, the HBSS with cells was removed using a large gauge needle. The solution was then overlaid on top of a 0/52/62.5/78 percent Percoll gradient and centrifuged at 700g for 30 minutes without acceleration or brake. Individual layers were removed by transfer pipette to individual tubes for washing. Wright’s staining (Wright’s Giemsa, Sigma-Aldrich WG16-500mL) was performed on 10-25µL of washed layers that were smeared and allowed to dry on slides before fixation in 100% methanol.

### Luminescent tumor growth tracking

Right-side midsections of mice were shaved and LLC1 cells (2×10^5^) were implanted subcutaneously in HBSS. Tumors were allowed to grow to an average of ∼200mm^3^ and then injected retro-orbitally with 1×10^6^ ST electroporated with the pAKlux2 plasmid. Mice were imaged for luminescence in the Small Animal Imaging Core at City of Hope using the Lago biophotonic imaging system (Spectral Instruments).

### Subcutaneous tumor growth and treatment

LLC1 cells were injected subcutaneously (2×10^5^ per mouse) in the right-side midsection of the mice. After 6 days of growth, treatment with shScr-ST or shIDO-ST began with three consecutive daily doses of 1×10^6^ cfu per mouse. On the first day of ST treatment, mice were concurrently given 200µg of anti-PD-1 (clone J43) antibody or IgG equivalent (BioXCell) and 75µg of anti-CTLA-4 (9H10) antibody or IgG equivalent (BioXCell). Treatments continued every three days until the end of the experiment (maximum tumor growth of 15 mm diameter for control groups). Tumor volumes were measured thrice weekly using manual calipers.

### Flow cytometry

One million live cells were counted using trypan blue and first stained with a fixable viability dye (eBiosciences, 65-0866-14) for 30 min at 4°C. Cells were washed in flow wash buffer (PBS with 0.05% sodium azide and 1% FBS) and stained with surface antibodies for 40 min at 4°C. Cells were washed in flow buffer and fixed in flow buffer plus 1% PFA before filtering through 40 μM mesh strainer/tube (BD Biosciences). Flow cytometry was performed on the BD FACSCelesta cytometer and data was analyzed using FlowJo Version 10 (Becton, Dickinson & Co.). For flow cytometry from spleens, spleens were removed and mashed in complete RPMI medium (10% FBS with glutamine and pen/strep) on a 70µm mesh strainer with a syringe plunger. Cells were washed in PBS and stained using the above protocol. For flow cytometry on tumor cells, tumors were excised and digested mechanically by mincing with a sterile scalpel before digestion in 1 mg/mL collagenase I (Sigma, C5138) plus 1% FBS for 1.5 h shaking at 200 rpm in a 37°C incubator. Dissociated tumor cells were spun at 450 g for 10 min and filtered through a 70 μM strainer. After washing, these cells were stained for flow cytometry following the above protocol. Flow cytometry antibodies used from BD biosciences: CD45-PerCPCy5.5 (550994), CD8-APC-R700 (564983), CD4-APC-H7 (560246), Ly6G-BV605 (563005), Ly6C-FITC (561085), CD11b-APC (561690), PD-1-BV421 (562584), CTLA-4-PE (553720), CD45-APC-R700 (565478), CD3-PerCPCy5.5 (551071), NK1.1-BV650 (564143), CD11c-BV605 (563057), CD45-BV650 (563410), CD4-BV786 (563331), CD25-PerCPcy5.5 (551071), MHCII-PerCPCy5.5 (562363), CD11c-APC-R700 (565871), IFN-gamma-APC (554413), CD27-BV450 (561245), andCD62L-BV605 (562720). Flow cytometry antibodies used from eBiosciences: F4/80-Super Bright 780 (78-4801-82), PD-L1-Super Bright 645 (64-5982-82), PD-L1-Super Bright 780 (78-5982-80), CD86-Super Bright 645 (64-0862-80), FoxP3-APC (17-5773-82), and Granzyme B-PE (12-8898-80). For intracellular staining against CTLA-4, cells were permeabilized using the BD Cytofix/Cytoperm Plus kit (555028). For intracellular staining against FoxP3, cells were permeabilized using the FoxP3/Transcription Factor staining buffer set from eBiosciences (00-5523-00).

### Statistics

All statistical analyses were done using the Prism software by GraphPad (V8). Unless otherwise noted in figure legends, statistics were obtained by performing a 2-way ANOVA followed by Tukey’s multiple comparisons test.

## RESULTS

### High-dose shIDO-ST treatment skews immunity towards a highly focused neutrophil response

Therapeutic shIDO-ST treatment has been shown to induce a predominantly cytotoxic neutrophil response that is directly involved in rapid clearance of tumor cells [22, 35]. To better assess the potential synergy between shIDO-ST and ICB treatment, we first utilized tumor-free C57BL/6 mice to circumvent mechanisms of suppression exerted by NSCLC tumors. Intravenous injection of therapeutic doses of shIDO-ST, or scrambled shRNA control (shScr-ST), resulted in an increase of neutrophils in the bloodstream upwards of ten times higher than in vehicle-injected mice (no ST) within 48 hours **(Figure 1A)**. By 72 hours, neutrophils in the bloodstream began to decrease in shScr-ST- and PBS-treated mice, but remained significantly higher (*p*<0.022) in shIDO-ST-treated mice. The percentage of Ly6G^+^ neutrophils was also significantly elevated (*p*<0.0007) in the spleens of mice administered shIDO-ST, compared to shScr-ST control treatment **(Figure 1B)**. Interestingly, phenotypic analysis of splenic neutrophils from shIDO-ST-treated mice revealed dramatically increased expression of professional APC markers, which included MHCII and CD80, compared to control-treated mice which may explain the additional adaptive immune response generated in previous models [22, 35]. These observations are in line with literature reporting the antigen presentation capabilities of certain neutrophil populations [38, 39].

**Figure 1.**
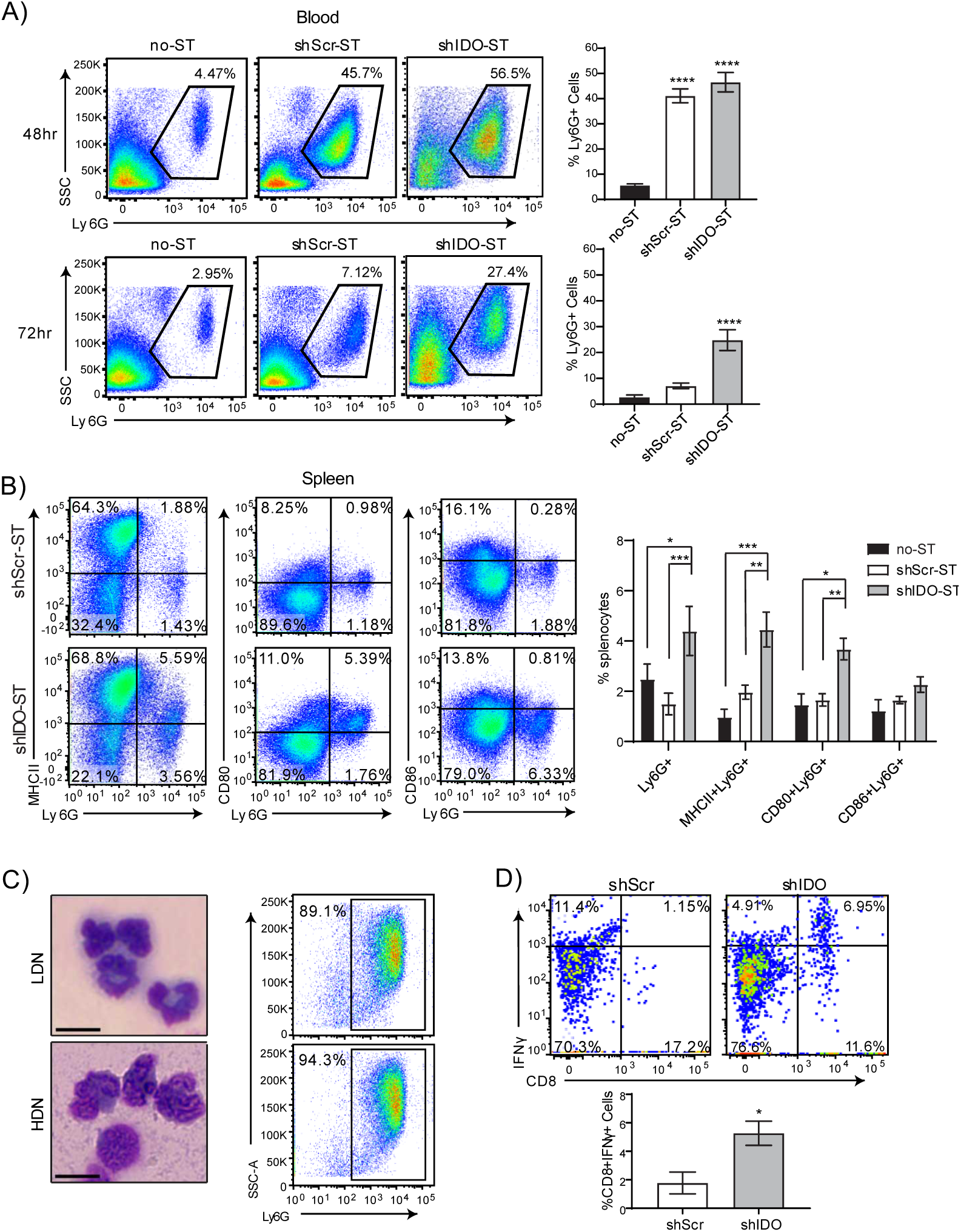
Neutrophil phenotypes after shIDO-ST treatment in tumor-free mice. **(A)** Flow cytometry showing neutrophil frequencies in blood by Ly6G staining out of CD45^+^ cells 48 or 72 hours after treatment with HBSS (no-ST), shScr-ST or shIDO-ST. Representative gated dot plots are shown. Bar graphs show quantification of flow cytometry results. n=4 mice per group. Statistics shown are compared to no-ST. **(B)** Flow cytometry showing double positivity for Ly6G and either MHCII, CD80, or CD86 for splenocytes 72 hours after treatment with shScr or shIDO-ST. Representative gated dot plots are shown. Bar graphs show quantification of flow cytometry results. n=4 mice per group. **(C)** Brightfield images (Wright’s stain) of cells isolated from the interfaces between 52 and 62.5% Percoll layers (LDN, low density neutrophils) and the interface between 62.5 and 78% Percoll layers (HDN, high density neutrophils). Scale bar = 10µm. Flow cytometry dot plots represent the purity of those fractions by Ly6G positivity. **(D)** Flow cytometry of activated CD8 T cells (CD8 and IFNγ double positive) out of total OTI splenocytes after 24 hours of *in vitro* co-incubation with SIINFEKL-loaded HDN from either shScr or shIDO-ST treated mice. Bar graph shows quantification of flow cytometry results. n=3 mice per group. Unpaired t-test. **(A-B, D)** *p<0.05, **p<0.01, ***p<0.001,****p<0.0001.

To determine the antigen-presenting capacity of neutrophils induced by shIDO-ST treatment, a Percoll gradient was used to isolate neutrophils for *in vitro* stimulation assays [40]. To obtain the highest yield of neutrophils for this experiment, shIDO-ST or shScr-ST were injected directly into the peritoneal cavity of mice, which reliably induces a large influx of neutrophils. Separation using 4-layer Percoll gradient enabled isolation of two populations of neutrophils, low- and high-density, with the higher density neutrophils showing extreme nuclear segmentation consistent with neutrophil activation **(Figure 1C)** [41, 42]. The predominance of low- and high-density cells isolated by Percoll gradient were Ly6G^+^ (≥90% and ≥94% purity, respectively). Purified neutrophil populations were then fixed and incubated with the ovalbumin synthetic peptide SIINFEKL, followed by incubation with splenocytes from OT-I mice transgenic for the SIINFEKL-specific T cell receptor [43]. Following co-incubation, splenocytes were analyzed for activated CD8^+^IFNγ^+^ T cells. In shIDO-ST-treated mice, the percentage of CD8^+^IFNγ^+^ T cells was significantly higher compared to shScr-ST-treated mice **(Figure 1D**, *p*<0.05), indicating an enhanced ability of shIDO-ST-induced neutrophils to augment SIINFEKL-specific T cell responses. As expected, these results were only observed in the high-density (activated) neutrophil fraction. Low-density neutrophils from shIDO-ST- or shScr-ST-treated mice did not significantly induce CD8^+^IFNγ^+^ responses **(Supplemental Fig 1A)**. These results are the first to report a potentially novel approach to generate functional hybrid APC-neutrophils, however, the immune response is highly skewed towards neutrophils, which may not be optimal for improving ICB treatment. Therefore, we next determined the effects of sub-therapeutic shIDO-ST doses in modifying the immune contexture.

### Low-dose shIDO-ST colonizes NSCLC tumors and does not induce cytotoxic neutrophils

In previous studies, high-dose shIDO-ST treatment (5×10^6^ cfu) induced transient intratumoral infiltration of cytotoxic neutrophils that mediated apoptosis of tumor-associated immune cells [22, 35, 44]. For these studies, we hypothesized that significantly reducing the dose of shIDO-ST might lead to a less exaggerated, cytotoxic neutrophil response so that other immune subsets, such as tumor-infiltrating T cells, might be preserved or augmented. We first confirmed that the ST vector, used to generate shIDO-ST, could colonize NSCLC tumors at a dose 5-fold lower than previously published (1×10^6^ cfu) [45, 46]. We engineered the ST vector to express the bioluminescent LUX reporter (LUX-ST) [47] in order to track movement *in vivo* using intravital imaging. We observed that intravenous injection of 1×10^6^ cfu LUX-ST [48] resulted in colonization of LLC1 tumors within 24 hours (**Figure 2A**). Bioluminescent signal in tumors peaked by day 3 and then began to decline, with measurable levels still detectable 7 days after LUX-ST injection. To ensure that 1×10^6^ cfu of shIDO-ST did not have immediate, cytotoxic anti-tumor activity as observed using high-dose shIDO-ST [35, 45, 49, 50], we treated LLC1 tumors with three daily doses of 1×10^6^ cfu shIDO-ST. This treatment regimen had no significant effect on tumor growth (**Figure 2B**) and did not change the frequency or APC status of tumor-associated neutrophils (**Figure 2C, D**). These results suggest that sub-therapeutic doses of shIDO-ST can effectively colonize LLC1 tumors without expanding a cytotoxic neutrophil response that might diminish other potential anti-tumor immune subsets.

**Figure 2.**
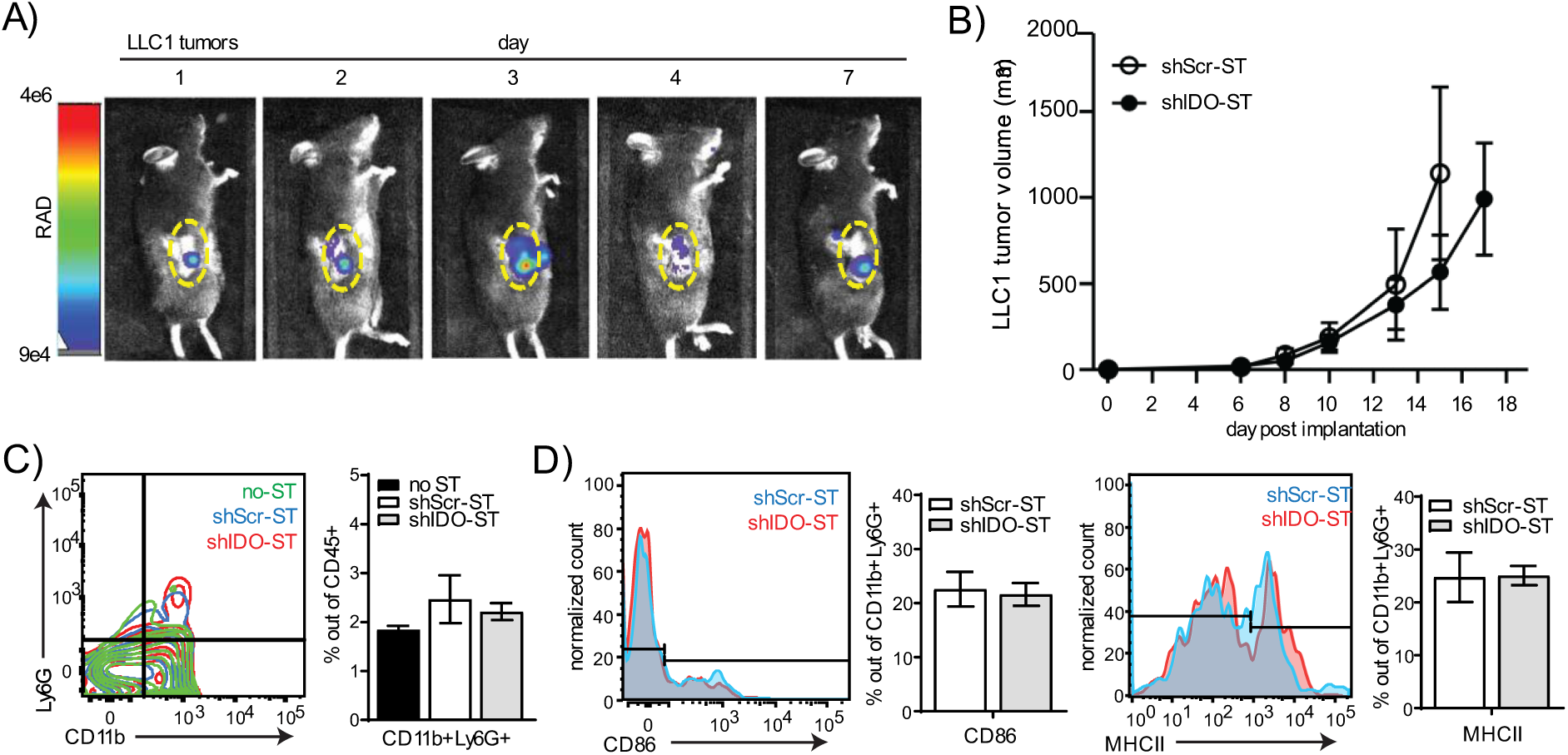
Colonization, tumor growth, and neutrophil phenotypes after sub-therapeutic shIDO-ST treatment. **(A)** Mice were implanted with LLC1 tumor cells subcutaneously and allowed to grow to an average volume of 200mm^3^. 1×10^6^ cfu of ST carrying a bacterial luciferase (LUX-ST) was injected intravenously and mice were imaged for bioluminescent signal on days 1, 2, 3, 4, and 7 post ST administration. Bioluminescent signal is shown in rainbow corresponding to radiance (RAD). Yellow dotted line borders the tumor. **(B)** Six days after implantation of LLC1 cells into mice, mice were treated with three consecutive, daily doses of 1×10^6^ cfu shScr or shIDO-ST. Tumor volumes were measured three times weekly until maximum allowed tumor growth was reached. n=4 mice per group. **(C)** Flow cytometry of neutrophil frequency (CD11b^+^Ly6G^+^) out of total CD45^+^ cells from tumors treated with HBSS (no-ST), shScr-ST or shIDO-ST. Tumors were processed 48 hours after the third treatment. Bar graph shows quantification of flow cytometry results. **(D)** Flow cytometry of antigen presentation molecules on neutrophils (CD86^+^ or MHCII^+^ out of CD11b^+^Ly6G^+^ cells). Bar graph shows quantification of flow cytometry results.

### Sub-therapeutic doses of shIDO-ST preserves intratumoral T cell frequency, increases antigen presentation potential and reduces Treg frequency

We next determined if sub-therapeutic shIDO-ST treatment allowed for changes in the immune landscape that could potentially augment ICB treatment. Using flow cytometry, we found that the frequencies of tumor-associated CD4^+^ and CD8^+^ T cells and myeloid-derived immune subsets 48 hours post-shIDO-ST treatment were comparable to PBS and shScr-ST control groups **(Figure 3A)**, suggesting that these populations are preserved in the absence of a cytotoxic neutrophil response [22]. Additionally, while low-dose shIDO-ST treatment did not expand hybrid APC-neutrophils (**Figure 2D**), we did observe increases in antigen presentation machinery and co-stimulatory markers on CD11c^+^ dendritic cell, CD11b^+^F4/80^+^ macrophage, and CD11b^+^Ly6C^+^ monocyte populations, which included increased CD86 and MHCII surface expression **(Figures 3B and 3C)**. Similar to tumor derived neutrophils (CD11b^+^Ly6G^+^), activated monocytes are difficult to distinguish from monocytic-MDSC. However, the increased expression of MHCII and CD86 are indicative of anti-tumor monocytes that may also be capable of antigen presentation to T cells similar to neutrophils [51, 52]. Interestingly, coincident with increases in the antigen-presenting potential of CD11c^+^ and CD11b^+^ immune subsets, sub-therapeutic shIDO-ST treatment also significantly decreased the frequency of intratumoral CD4^+^CD25^+^FoxP3^+^ Tregs (p=.0475) **(Figure 3D)**. These results confirm that low-dose shIDO-ST treatment effectively alters the LLC1 tumor microenvironment by increasing antigen presentation potential and decreasing the presence of suppressive Tregs.

**Figure 3.**
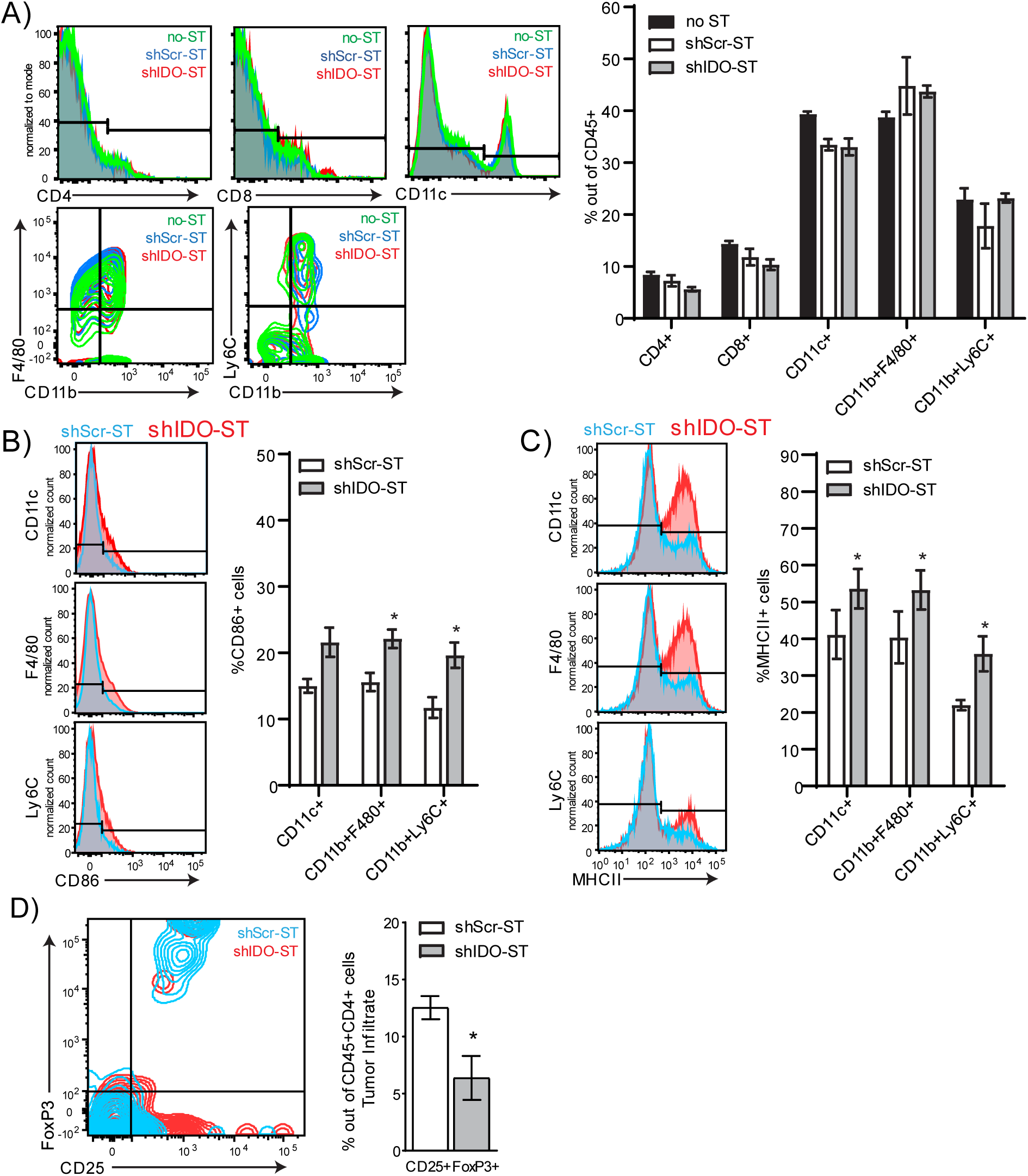
Frequency and phenotypes of tumor-infiltrating immune cells after shIDO-ST treatment. Mice with LLC1 tumors (average 80mm^3^) were treated with three consecutive doses of 1×10^6^ cfu of shScr-ST or shIDO-ST and tumors were processed for flow cytometry 48 hours after the third treatment. n=3 mice per group. **(A)** Frequencies of CD4^+^, CD8^+^, CD11c^+^, CD11b^+^F4/80^+^, and CD11b^+^Ly6C^+^ cells out of CD45^+^ tumor-infiltrating cells. **(B)** Frequency of CD86^+^ cells out of myeloid cell types (CD11c^+^, CD11b^+^F4/80^+^, and CD11b^+^Ly6C^+^ cells). Bar graphs show quantification of flow cytometry results. **(C)** Frequency of MHCII^+^ cells out of myeloid cell types (CD11c^+^, CD11b^+^F4/80^+^, and CD11b^+^Ly6C^+^ cells). Bar graphs show quantification of flow cytometry results. **(D)** Frequency of Tregs (CD25^+^FoxP3^+^ out of CD45^+^CD4^+^ cells). Bar graph shows quantification of flow cytometry results. Unpaired t-test. **(A-D)** *p<0.05, **p<0.01, ***p<0.001, ****p<0.0001.

### Sub-therapeutic shIDO-ST treatment suppresses checkpoint protein expression in splenic immune subsets

Besides interactions within the tumor, interactions between splenic T cells and suppressive immune subsets or those expressing checkpoint proteins may inhibit T cell priming against tumor antigens in the spleen [53], or mobilization of naïve T cells from the spleen to the tumor for priming there [54]. Splenic neutrophils, monocytes, dendritic cells, and macrophages show no changes in expression of the antigen presentation molecules CD86 and MHCII (**Supplemental Fig 2A-D**). However, total CD45^+^ cells are significantly increased in shIDO-ST-treated mice compared to shScr-ST-treated mice (p<0.0001) **(Supplemental Fig 2E)**, possibly representing an increase in B cells or another subset, or the combination of small increases in each cell type, as no specific cell type assayed was significantly increased **(Supplemental Fig 2F)**. Splenic Tregs were unchanged by shIDO-ST treatment **(Supplemental Fig 2G)**. However, shIDO-ST treatment was capable of decreasing expression of the checkpoint molecules PD-L1, PD-1, and CTLA-4 on certain cell types. PD-L1 surface expression on CD4^+^ and CD8^+^ T cells, CD11c^+^cells, macrophages, and NK cells is decreased significantly in shIDO-ST treated mice compared to shScr-ST treated mice **(Fig 4A)**. In addition, shIDO-ST treatment significantly decreases surface expression of the PD-1 receptor on CD4^+^ and CD8^+^ T cells, monocytes, and NK cells **(Fig 4B)**. Furthermore, shIDO-ST treatment significantly decreases CTLA-4 expression on CD4^+^ and CD8 T^+^ cells as well as NK cells **(Fig 4C)**. PD-1 surface expression on T cells correlates with activation, however high expression of PD-1 has been correlated with T cell anergy [55], similar findings have been published concerning PD-L1 and CTLA-4 on various immune subsets [56, 57]. Decreased expression of all three checkpoint proteins has the potential to globally decrease suppression, creating a TME more conducive to checkpoint blockade with both anti-PD-1 and anti-CTLA-4 antibodies.

**Figure 4.**
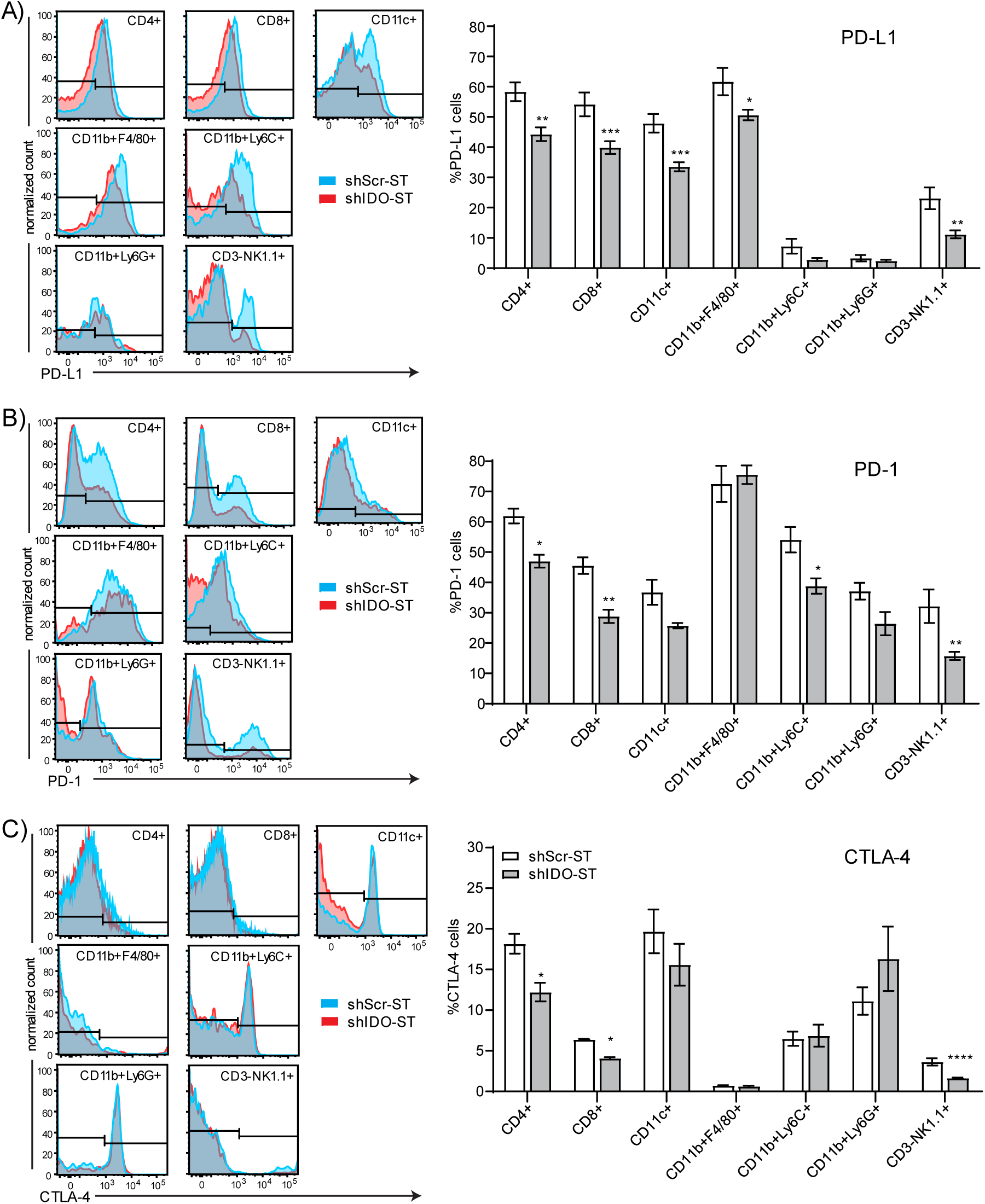
Checkpoint protein positivity of splenocytes by immune cell type after shIDO-ST treatment. Mice with LLC1 tumors (average 80mm^3^) were treated with three consecutive doses of 1×10^6^ cfu of shScr-ST or shIDO-ST and spleens were processed for flow cytometry 48 hours after the third treatment. n=3 mice per group. Cells were gated by CD45 positivity and then by cell type (CD4^+^, CD8^+^, CD11c^+^, CD11b^+^F4/80^+^, CD11b^+^Ly6C^+^, CD11b^+^Ly6G^+^, and CD3^-^NK1.1^+^). Flow cytometry is represented by histogram overlay followed by bar graph quantification of histogram results. **(A)** Frequency of PD-L1^+^ cells by immune cell type. **(B)** Frequency of PD-1^+^ cells by immune cell type. Frequency of CTLA-4^+^ cells by immune cell type. **(A-C)** *p<0.05, **p<0.01, ***p<0.001, ****p<0.0001.

### ShIDO-ST pre-treatment augments ICB efficacy and is associated with enhanced immune infiltration

Despite increased neutrophil infiltration and monocyte expression of APC-like markers in the tumor, as well as decreased numbers of suppressive Tregs in the tumor and decreased expression of checkpoint molecules in the spleen, shIDO-ST treatment is insufficient to activate intratumoral T cells. The intracellular expression of IFNγ in either CD4^+^ **(Supplemental Fig 3A)** or CD8^+^ T cells **(Supplemental Fig 3B)** remains unchanged between shScr-ST and shIDO-ST-treated groups, which may explain the lack of tumor control by subtherapeutic doses of shIDO-ST alone **(Fig 2B)**. However, the addition of anti-PD-1 and anti-CTLA-4 antibodies to shIDO-ST treatment (shIDO-ST+ICB) resulted in significant tumor control compared to shScr-ST+IgG or shIDO-ST+IgG as well as shScr-ST+ICB. Beginning on day 15, and continuing through the end of the study, the growth of shIDO-ST+ICB treated tumors was significantly reduced compared to all other groups: shScr-ST+IgG (p=0.0007), shIDO+IgG (p=0.0020) and shScr-ST+ICB (p=0.0020). Tumor growth in control groups did not differ significantly throughout the study **(Fig 5A)**. These data indicate that only shIDO-ST is capable of priming the TME for ICB treatment as tumors treated with shScr-ST+ICB continue to grow aggressively. To determine the cause of tumor control in the shIDO-ST+ICB treatment group, flow cytometry was used to quantify the numbers of CD45^+^ infiltrating immune cells in both the tumor and spleen. Overall CD45^+^ cells in the tumor increased significantly in the shIDO-ST+ICB group compared to shScr-ST+ICB or shIDO-ST groups (p<0.0001 for all groups) **(Fig 5B)**. Total CD45^+^ cells in the spleen remained unchanged between groups **(Fig 5C)**. A more in-depth analysis of specific immune subsets, however, revealed changes in the infiltration of multiple cell types in both the tumor and spleen. The shIDO-ST+ICB treated tumors have significantly higher numbers of tumor-infiltrating CD4^+^ and CD8^+^ T cells compared to all other treatment groups (p<0.0001), as well as significantly higher numbers of CD11c^+^ cells (p **(Fig 6A)**. Tumor-infiltrating CD11b^+^F4/80^+^ cells remained significantly lower in shIDO-ST+IgG and shIDO-ST+ICB groups compared to shScr-ST groups indicating that this effect is due to shIDO-ST treatment alone, whereas ICB alone decreased CD11b^+^Ly6C^+^ in both shScr and shIDO-ST groups. Tumor-infiltrating NK cells were significantly decreased only in the shIDO-ST+ICB treatment group **(Fig 6A)**. ICB treatment in both shScr-ST and shIDO-ST groups increased Tregs out of total CD4^+^ T cells consistent with reports in human tumor treatment [58, 59]; however, shIDO-ST treatment in combination with ICB prevented the level of Treg increase seen in the shScr-ST+ICB treatment group **(Fig 6B)**. These increased Treg after ICB treatment may explain the trend in increased tumor growth in the shScr-St+ICB treated group. CD4^+^ and CD8^+^ T cells were analyzed for expression of activation markers including CD27 (which promotes T cell survival and enhances priming) [60], CD62L (which is rapidly lost from T cells after specific antigen stimulation) [61], granzyme B (a cytotoxic factor) [62], Interferon gamma (IFNγ, a multi-cell immune modulator secreted by activated T cells) [63], and NK1.1 (in conjunction with CD3 positivity marks Natural Killer T cells (NKT)) [64]. Out of CD4^+^ and CD8^+^ cells in tumors, most markers remained unchanged between shScr-ST+ICB and shIDO-ST+ICB groups except for NK1.1, which was significantly decreased in shIDO-ST+ICB treated tumors compared to shScr-ST+ICB treated tumors (CD4^+^ p=0.0059, CD8^+^ p=0.0027) **(Fig 6C)**. However, due to the overall increase in CD4^+^ and CD8^+^ T cells in tumors, this denotes a greater amount of activated T cells present in tumors after shIDO-ST+ICB treatment compared to other groups.

**Figure 5.**
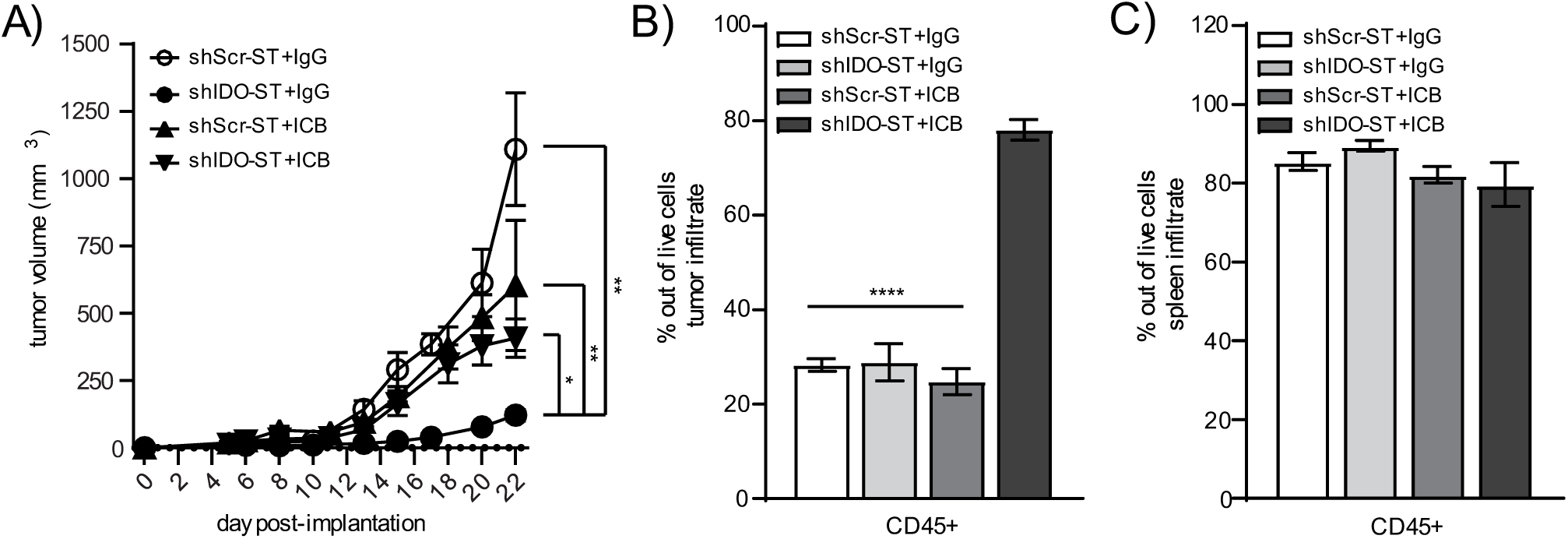
Tumor growth and total immune infiltrate after combination treatment with shScr or shIDO-ST and antibodies targeting PD-1 and CTLA-4. Mice with palpable LLC1 tumors were treated with three consecutive doses of 1×10^6^ cfu of shScr or shIDO-ST combined with immune checkpoint blockade (ICB, PD-1 and CTLA-4 antibodies) or IgG equivalent. PD-1 was administered at a dose of 200µg and CTLA-4 was administered at a dose of 75µg every three days until most groups reached maximum tumor growth (15cm diameter). n=4-10 mice per group. **(A)** Tumors were measured 3 times weekly and volume in mm^3^ is presented as a tumor growth curve. Statistical significance between shIDO-ST+ICB and shIDO-ST+IgG is shown for days 17, 20, 22, and 24. **(B)** Tumors and **(C)** spleens were excised 48 hours after the third ST treatment (24 hours after the second ICB or IgG treatment) and processed for flow cytometry. Frequencies of infiltrating CD45^+^ immune cells are shown in bar graphs.

**Figure 6.**
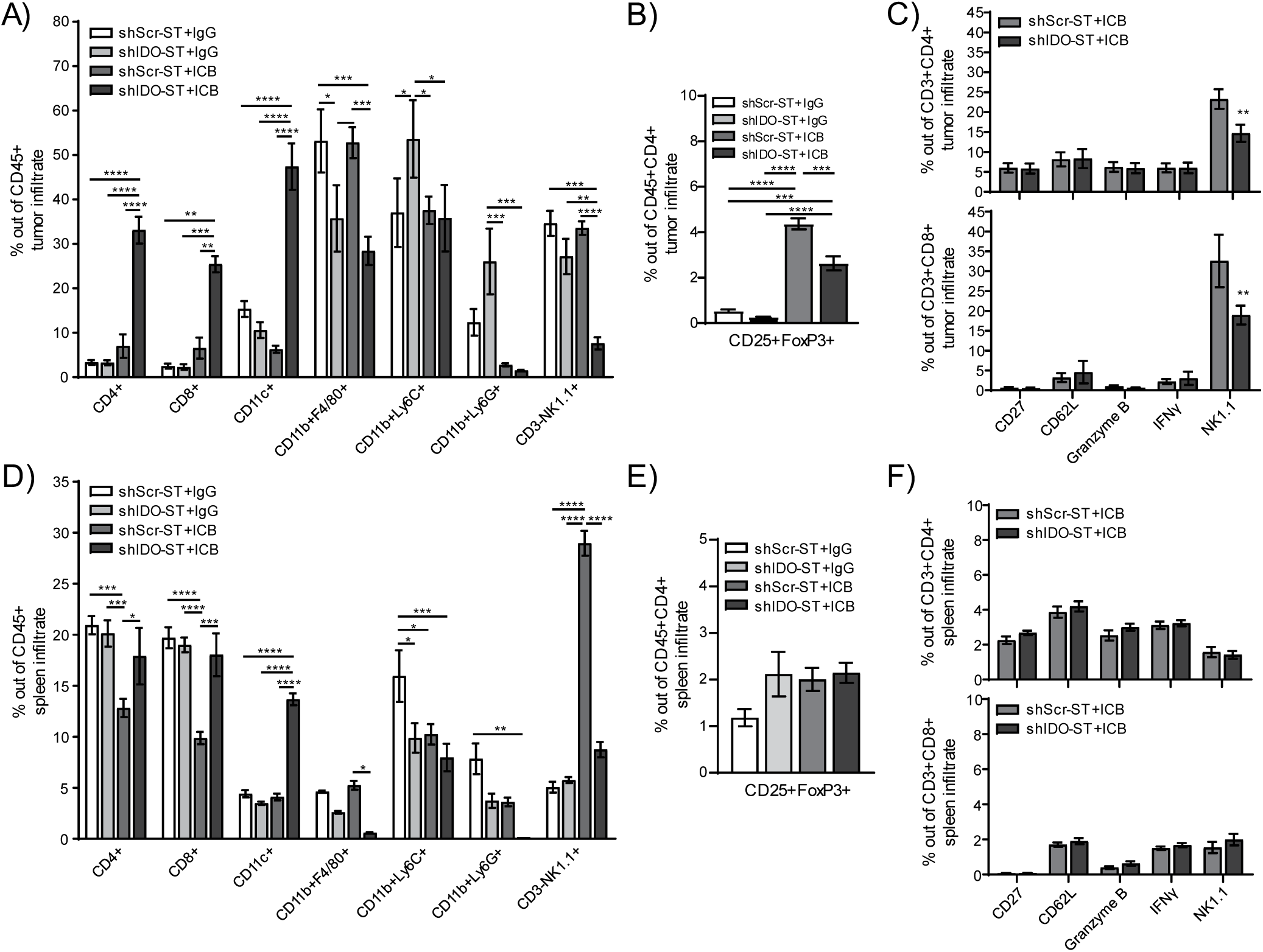
Frequency and phenotypes of tumor- and spleen-infiltrating immune cells after combination treatment with shScr or shIDO-ST and ICB. Mice with palpable LLC1 tumors were treated with three consecutive doses of 1×10^6^ cfu of shScr or shIDO-ST combined with immune checkpoint blockade (ICB, PD-1 and CTLA-4 antibodies) or IgG equivalent. Every three days, PD-1 was administered at a dose of 200µg and CTLA-4 was administered at a dose of 75µg until the end of the study. n=3-4 mice per group. **(A)** Tumors were excised 48 hours after the third ST treatment (24 hours after the second ICB or IgG treatment) and analyzed by flow cytometry for frequency of infiltrating immune cells out of the total CD45^+^ fraction. **(B)** Tumors were also analyzed by flow cytometry for the frequency of Tregs (CD25^+^FoxP3^+^ cells out of CD45^+^CD4^+^ cells). **(C)** T cells from tumors (CD3^+^CD4^+^ or CD3^+^CD8^+^) were analyzed by flow cytometry for activation though various markers (CD27, CD62L, granzyme B, IFNγ, and NK1.1). **(D)** Spleens were excised 48 hours after the third ST treatment (24 hours after the second ICB or IgG treatment) and analyzed by flow cytometry for frequency of infiltrating immune cells out of the total CD45^+^ fraction. **(E)** Spleens were also analyzed by flow cytometry for the frequency of Tregs (CD25^+^FoxP3^+^ cells out of CD45^+^CD4^+^ cells). **(F)** T cells from spleens (CD3^+^CD4^+^ or CD3^+^CD8^+^) were analyzed by flow cytometry for activation using various markers (CD27, CD62L, granzyme B, IFNγ, and NK1.1).

In the spleen, ICB treatment combined with shScr-ST decreases CD4^+^ and CD8^+^ T cells compared to IgG combined with either shScr-ST or shIDO-ST treatment; however, the combination of ICB with shIDO-ST prevents this decrease. In addition, shIDO-ST combined with ICB treatment alone increases CD11c^+^ cells and decreases CD11b^+^F4/80^+^ macrophages and CD11b^+^Ly6G^+^ neutrophils. Interestingly, shScr-ST with ICB increases the presence of CD3^-^NK1.1^+^ cells significantly in the spleen while shIDO-ST plus ICB combination prevents this increase **(Fig 6D)**. Tregs remained unchanged in the spleen **(Fig 6E)**, and similar to results seen in the tumor, activation markers for T cells also remained unchanged **(Fig 6F)**.

## DISCUSSION

Known mechanisms of ICB resistance include the presence of immune-suppressive cell types, such as Tregs and MDSCs, as well as mechanisms that preserve T cell tolerance despite ICB. Secretion of cytokines that maintain these mechanisms of resistance can come from tumor cells as well as many immune cell types, indicating that global changes in the TME as well as antigenic re-education of T cells may be necessary for tumor-infiltrating T cells to be fully activated by ICB. In this study we explore the ability of shIDO-ST treatment [22, 35, 44] to affect the immune landscape both in a tumor-free setting and using a syngeneic lung cancer model in mice. In the absence of a tumor, intravenous injection of mice with high-dose shIDO-ST dramatically increases neutrophils in the blood and spleen compared to mice injected with shScr-ST. Neutrophils present in the spleen after shIDO-ST injection showed increased expression of markers consistent with professional APCs, and isolated neutrophils from shIDO-ST injected mice activated IFNγ production in T cells via antigen-specific MHC class I presentation. These results are consistent with anti-tumor, APC-like neutrophils found in patients with lung cancer [30, 31, 65].

Gene expression studies of auto-reactive T cells have revealed that T cell dysfunction occurs in phases and that these various T cell sub-populations differ in their response to checkpoint blockade [66-68]. Fully tolerized, self T cells—which share a very similar transcriptional profile to fixed, dysfunctional, tumor-specific T cells [69]—are completely resistant to single target checkpoint blockade treatment as well as the combination of anti-CTLA-4 and anti-PD-1. Remarkably, despite down-regulation of genes related to antigen response, re-introduction of auto-antigens has been shown to reverse tolerance and sensitize T cells to checkpoint blockade [66]. These data indicate that re-establishment of the full range of immune cell activity, or induction of novel immune functions, may be necessary for enhancing efficacy of ICB treatment. In LLC1 tumor-bearing mice, low-dose shIDO-ST treatment preserves T cell populations and monocytes experience an increase in CD86 and MHCII expression indicating that these myeloid cells, similar to neutrophils in the tumor-free model, may have greater ability to present tumor-specific antigen to tolerized T cells. Tumor Tregs are also significantly decreased by shIDO-ST treatment. In the spleen, expression of PD-1, PD-L1, and CTLA-4 are decreased in multiple immune subsets. These data indicate that shIDO-ST treatment is capable of altering the TME, possibly resulting in decreased suppression and increased antigen presentation which would increase the efficacy of ICB.

The addition of antibodies blocking PD-1 and CTLA-4 to shIDO-ST treatment results in increased tumor control compared to shIDO-ST with IgG or shScr-ST with ICB treatment. This increased control may be attributed to an overall increase in the CD45^+^ fraction of tumor cells including CD4^+^ and CD8^+^ T cells, as well as CD11c^+^ cells. Notably downregulated cells types include CD11b^+^F4/80^+^ macrophages, CD11b^+^Ly6G^+^ neutrophils, and CD3^-^NK1.1^+^ NK cells, possibly indicating that these cell types are suppressive in the LLC1 model. Percentages of activated tumor-infiltrating T cells remained unchanged between shScr-ST+ICB and shIDO-ST+ICB groups, but the overall increase in these T cells in the shIDO-ST+ICB group indicates greater numbers of activated T cells in the tumor overall. Interestingly, NK1.1 expression on CD3^+^CD4^+^ or CD8^+^ T cells was decreased in shIDO-ST+ICB treated tumors compared to shScr-ST+ICB treated tumors, indicating fewer NKT. These cell types have previously been shown to either promote or hinder anti-tumor immunity depending on their cytokine production profile, with type I NKTs producing IFNγ to recruit NK cells and activate DCs [70], whereas Type II NKTs produce IL-13 and TGF-β that suppress anti-tumor immunity [71, 72]. In tumors, suppressive type II NKTs may dominate due to the presence of Tregs [73], and ICB may increase this population due to the increased Tregs seen in Fig 6C. However, shIDO-ST+ICB treatment prevented the increase in this suppressive population despite the Treg increase.

Taken together, these data indicate that changes occurring in certain cell types after shIDO treatment alone prime the TME for treatment with ICB. One study found that T cells newly introduced to antigen were capable of responding favorably to ICB, but T cells tolerized to self-antigen or tumor-antigen did not respond to ICB alone [66]. However, re-introduction of antigen synergized with ICB to produce functional T cells capable of eliminating tumor cells expressing self- or tumor-antigens [66]. Our data showing increased expression of antigen presentation machinery on the surface of monocytes and decreased Treg abundance in shIDO-ST treated mice may have increased antigen presentation to T cells overall, thus priming the TME for ICB treatment which only increased T cell infiltration in combination with shIDO-ST. In addition, decreasing immune checkpoint proteins on the surfaces of multiple cell types, as we have shown occurs after shIDO-ST treatment in splenocytes, may aid in increasing antigen presentation. One study showed that blocking PD-L1 with antibody on MDSCs causes a change in cytokine secretion, namely downregulation of IL-6 and IL-10 [29], which could have profound effects on the TME.

The link between IDO expression in the TME and antigen presentation is largely unknown, but some mechanisms have been described concerning the effects of agonists of the aryl hydrocarbon receptor (AhR) similar to kynurenine (the product of tryptophan conversion by IDO). In addition, to the deleterious effects of kynurenine on T cells [74, 75], agonists of the AhR have been shown to regulate the immunogenicity of other cell types including APCs. Administration of the AhR agonist 2-(1’H-indole-3’-carbonyl)-thiazole-4-carboxylic acid methyl ester (ITE) to mice prevented the development of experimental autoimmune encephalomyelitis through tolerization of dendritic cells after decreasing their expression of CD86 while increasing secretion of anti-inflammatory cytokines such as IL-10 and TGF-β [76]. In vitro, ITE-treated bone marrow-derived DCs were unable to activate T cells in co-culture, and passive transfer of ITE-treated DCs were alone able to significantly inhibit experimental autoimmune encephalomyelitis [76]. In an allergy model, DCs isolated from the lungs of AhR^-/-^ mice were able to stimulate greater T cell responses to specific antigen *in vitro* and showed higher expression of CD86 and MHCII compared with wildtype DCs [77]. These data indicate that decreased production of AhR agonists such as kynurenine after IDO inhibition or knockdown can directly affect the antigen presentation capabilities of professional APCs such as DCs. If these pathways overlap in other immune cell types such as neutrophils or monocytes, then IDO knockdown, via shIDO-ST, may be directly responsible for upregulation of their ability to present antigen, and the overall effects of IDO knockdown may account for ICB synergy due to increased antigen presentation through multiple cell types in the LLC1 lung cancer model.

The ability of shIDO-ST to decrease the suppressive immune phenotype of the TME in the LLC1 NSCLC model represents a promising mechanism to increase the action of ICB treatments in human NSCLC patients. The ability to increase the number of patients that will respond effectively and durably to ICB treatments will fill an unmet need within the cancer care community and may represent an avenue for increasing the efficacy of ICB in other tumor types that are as yet unresponsive.

## ACKNOWLEDGMENTS

This study was funded by an NCCN Young Investigator Award and COH Shared Resources grant (both to ERM). The COH Comprehensive Cancer Center is supported by P30 CA033572.

## REFERENCES

1. Kargl J, Busch SE, Yang GHY, Kim K-H, Hanke ML, Metz HE, Hubbard JJ, Lee SM, Madtes DK, McIntosh MW et al: Neutrophils dominate the immune cell composition in non-small cell lung cancer. Nature Communications 2017, 8(1):14381.

2. Stankovic B, Bjørhovde HAK, Skarshaug R, Aamodt H, Frafjord A, Müller E, Hammarström C, Beraki K, Bækkevold ES, Woldbæk PR et al: Immune Cell Composition in Human Non-small Cell Lung Cancer. Front Immunol 2019, 9(3101).

3. Prado-Garcia H, Romero-Garcia S, Morales-Fuentes J, Aguilar-Cazares D, Lopez-Gonzalez JS: Activation-induced cell death of memory CD8+ T cells from pleural effusion of lung cancer patients is mediated by the type II Fas-induced apoptotic pathway. Cancer Immunol, Immunother 2012, 61(7):1065–1080.

4. Prado-Garcia H, Aguilar-Cazares D, Flores-Vergara H, Mandoki JJ, Lopez-Gonzalez JS: Effector, memory and naïve CD8+ T cells in peripheral blood and pleural effusion from lung adenocarcinoma patients. Lung Cancer 2005, 47(3):361–371.

5. Shi L, Chen S, Yang L, Li Y: The role of PD-1 and PD-L1 in T-cell immune suppression in patients with hematological malignancies. Journal of hematology & oncology 2013, 6(1):74.

6. Pawelczyk K, Piotrowska A, Ciesielska U, Jablonska K, Gletzel-Plucinska N, Grzegrzolka J, Podhorska-Okolow M, Dziegiel P, Nowinska K: Role of PD-L1 Expression in Non-Small Cell Lung Cancer and Their Prognostic Significance according to Clinicopathological Factors and Diagnostic Markers. Int J Mol Sci 2019, 20(4):824.

7. Zhang N, Tu J, Wang X, Chu Q: Programmed cell death-1/programmed cell death ligand-1 checkpoint inhibitors: differences in mechanism of action. Immunotherapy 2019, 11(5):429–441.

8. Rizvi NA, Hellmann MD, Snyder A, Kvistborg P, Makarov V, Havel JJ, Lee W, Yuan J, Wong P, Ho TS et al: Cancer immunology. Mutational landscape determines sensitivity to PD-1 blockade in non-small cell lung cancer. Science 2015, 348(6230):124–128.

9. McGranahan N, Furness AJ, Rosenthal R, Ramskov S, Lyngaa R, Saini SK, Jamal-Hanjani M, Wilson GA, Birkbak NJ, Hiley CT et al: Clonal neoantigens elicit T cell immunoreactivity and sensitivity to immune checkpoint blockade. Science 2016, 351(6280):1463–1469.

10. Somasundaram A, Burns TF: Pembrolizumab in the treatment of metastatic non-small-cell lung cancer: patient selection and perspectives. Lung Cancer (Auckl) 2017, 8:1–11.

11. Steendam CM, Dammeijer F, Aerts J, Cornelissen R: Immunotherapeutic strategies in non-small-cell lung cancer: the present and the future. Immunotherapy 2017, 9(6):507–520.

12. Seetharamu N, Preeshagul IR, Sullivan KM: New PD-L1 inhibitors in non-small cell lung cancer - impact of atezolizumab. Lung Cancer (Auckl) 2017, 8:67–78.

13. Krummel MF, Allison JP: CD28 and CTLA-4 have opposing effects on the response of T cells to stimulation. J Exp Med 1995, 182(2):459–465.

14. Lynch TJ, Bondarenko IN, Luft A, Serwatowski P, Barlesi F, Chacko RT, Sebastian M, Siegel J, Cuillerot J, Reck M: Phase II trial of ipilimumab (IPI) and paclitaxel/carboplatin (P/C) in first-line stage IIIb/IV non-small cell lung cancer (NSCLC). J Clin Oncol 2010, 28(15_suppl):7531–7531.

15. Hellmann MD, Paz-Ares L, Bernabe Caro R, Zurawski B, Kim S-W, Carcereny Costa E, Park K, Alexandru A, Lupinacci L, de la Mora Jimenez E et al: Nivolumab plus Ipilimumab in Advanced Non–Small-Cell Lung Cancer. New Engl J Med 2019, 381(21):2020–2031.

16. Dudnik E, Moskovitz M, Daher S, Shamai S, Hanovich E, Grubstein A, Shochat T, Wollner M, Bar J, Merimsky O et al: Effectiveness and safety of nivolumab in advanced non-small cell lung cancer: The real-life data. Lung cancer 2017.

17. Carbone DP, Reck M, Paz-Ares L, Creelan B, Horn L, Steins M, Felip E, van den Heuvel MM, Ciuleanu T-E, Badin F et al: First-Line Nivolumab in Stage IV or Recurrent Non-Small-Cell Lung Cancer. The New England journal of medicine 2017, 376(25):2415–2426.

18. Lin GN, Peng JW, Liu PP, Liu DY, Xiao JJ, Chen XQ: Elevated neutrophil-to-lymphocyte ratio predicts poor outcome in patients with advanced non-small-cell lung cancer receiving first-line gefitinib or erlotinib treatment. Asia-Pacific journal of clinical oncology 2017, 13(5):e189–e194.

19. Jiang L, Zhao Z, Jiang S, Lin Y, Yang H, Xie Z, Lin Y, Long H: Immunological markers predict the prognosis of patients with squamous non-small cell lung cancer. Immunologic research 2015, 62(3):316–324.

20. Ngiow SF, Young A, Jacquelot N, Yamazaki T, Enot D, Zitvogel L, Smyth MJ: A Threshold Level of Intratumor CD8<sup>+</sup> T-cell PD1 Expression Dictates Therapeutic Response to Anti-PD1. Cancer Res 2015, 75(18):3800–3811.

21. Paluskievicz CM, Cao X, Abdi R, Zheng P, Liu Y, Bromberg JS: T Regulatory Cells and Priming the Suppressive Tumor Microenvironment. Front Immunol 2019, 10(2453).

22. Blache CA, Manuel ER, Kaltcheva TI, Wong AN, Ellenhorn JDI, Blazar BR, Diamond DJ: Systemic Delivery of Salmonella Typhimurium Transformed with IDO shRNA Enhances Intratumoral Vector Colonization and Suppresses Tumor Growth. Cancer Res 2012, 72(24):6447–6456.

23. Corzo CA, Cotter MJ, Cheng P, Cheng F, Kusmartsev S, Sotomayor E, Padhya T, McCaffrey TV, McCaffrey JC, Gabrilovich DI: Mechanism regulating reactive oxygen species in tumor induced myeloid-derived suppressor cells: MDSC and ROS in cancer. J Immunol 2009, 182(9):5693–5701.

24. Raber P, Ochoa AC, Rodríguez PC: Metabolism of L-Arginine by Myeloid-Derived Suppressor Cells in Cancer: Mechanisms of T cell suppression and Therapeutic Perspectives. Immunol Invest 2012, 41(6-7):614–634.

25. Mbongue JC, Nicholas DA, Torrez TW, Kim N-S, Firek AF, Langridge WHR: The Role of Indoleamine 2, 3-Dioxygenase in Immune Suppression and Autoimmunity. Vaccines 2015, 3(3):703–729.

26. Li H, Han Y, Guo Q, Zhang M, Cao X: Cancer-Expanded Myeloid-Derived Suppressor Cells Induce Anergy of NK Cells through Membrane-Bound TGF-β1. The Journal of Immunology 2009, 182(1):240–249.

27. Mittal SK, Roche PA: Suppression of antigen presentation by IL-10. Curr Opin Immunol 2015, 34:22–27.

28. Park M-J, Lee S-H, Kim E-K, Lee E-J, Baek J-A, Park S-H, Kwok S-K, Cho M-L: Interleukin-10 produced by myeloid-derived suppressor cells is critical for the induction of Tregs and attenuation of rheumatoid inflammation in mice. Sci Rep 2018, 8(1):3753.

29. Noman MZ, Desantis G, Janji B, Hasmim M, Karray S, Dessen P, Bronte V, Chouaib S: PD-L1 is a novel direct target of HIF-1α, and its blockade under hypoxia enhanced MDSC-mediated T cell activation. J Exp Med 2014, 211(5):781–790.

30. Singhal S, Bhojnagarwala PS, O’Brien S, Moon EK, Garfall AL, Rao AS, Quatromoni JG, Stephen TL, Litzky L, Deshpande C et al: Origin and Role of a Subset of Tumor-Associated Neutrophils with Antigen-Presenting Cell Features in Early-Stage Human Lung Cancer. Cancer Cell 2016, 30(1):120–135.

31. Saha S, Biswas SK: Tumor-Associated Neutrophils Show Phenotypic and Functional Divergence in Human Lung Cancer. Cancer Cell 2016, 30(1):11–13.

32. Granot Z, Henke E, Comen EA, King TA, Norton L, Benezra R: Tumor entrained neutrophils inhibit seeding in the premetastatic lung. Cancer cell 2011, 20(3):300–314.

33. Eruslanov EB: Phenotype and function of tumor-associated neutrophils and their subsets in early-stage human lung cancer. Cancer Immunol Immunother 2017, 66(8):997–1006.

34. D’Alincourt Salazar M, Manuel ER, Tsai W, D’Apuzzo M, Goldstein L, Blazar BR, Diamond DJ: Evaluation of innate and adaptive immunity contributing to the antitumor effects of PD1 blockade in an orthotopic murine model of pancreatic cancer. Oncoimmunology 2016, 5(6):e1160184.

35. Manuel ER, Chen J, D’Apuzzo M, Lampa MG, Kaltcheva TI, Thompson CB, Ludwig T, Chung V, Diamond DJ: Salmonella-Based Therapy Targeting Indoleamine 2,3-Dioxygenase Coupled with Enzymatic Depletion of Tumor Hyaluronan Induces Complete Regression of Aggressive Pancreatic Tumors. Cancer Immunol Res 2015, 3(9):1096–1107.

36. Chen J, Diamond DJ, Manuel ER: Developing Effective Salmonella-based Approaches to Treat Pancreatic Cancer. Pancreat Disord Ther 2016, 6(1):1–2.

37. Ebelt ND, Manuel ER: Utilizing Salmonella to treat solid malignancies. Journal of surgical oncology 2017, 116(1):75–82.

38. Matsushima H, Geng S, Lu R, Okamoto T, Yao Y, Mayuzumi N, Kotol PF, Chojnacki BJ, Miyazaki T, Gallo RL et al: Neutrophil differentiation into a unique hybrid population exhibiting dual phenotype and functionality of neutrophils and dendritic cells. Blood 2013, 121(10):1677–1689.

39. Singhal S, Bhojnagarwala Pratik S, O’Brien S, Moon Edmund K, Garfall Alfred L, Rao Abhishek S, Quatromoni Jon G, Stephen Tom L, Litzky L, Deshpande C et al: Origin and Role of a Subset of Tumor-Associated Neutrophils with Antigen-Presenting Cell Features in Early-Stage Human Lung Cancer. Cancer Cell 2016, 30(1):120–135.

40. Swamydas M, Luo Y, Dorf ME, Lionakis MS: Isolation of Mouse Neutrophils. Curr Protoc Immunol 2015, 110:3.20.21-23.20.15.

41. Sagiv Jitka Y, Michaeli J, Assi S, Mishalian I, Kisos H, Levy L, Damti P, Lumbroso D, Polyansky L, Sionov Ronit V et al: Phenotypic Diversity and Plasticity in Circulating Neutrophil Subpopulations in Cancer. Cell Rep 2015, 10(4):562–573.

42. Aulakh GK, Balachandran Y, Liu L, Singh B: Angiostatin inhibits activation and migration of neutrophils. Cell Tissue Res 2014, 355(2):375–396.

43. Hogquist KA, Jameson SC, Heath WR, Howard JL, Bevan MJ, Carbone FR: T cell receptor antagonist peptides induce positive selection. Cell 1994, 76(1):17–27.

44. Phan T, Nguyen VH, D’Alincourt MS, Manuel ER, Kaltcheva T, Tsai W, Blazar BR, Diamond DJ, Melstrom LG: Salmonella-mediated therapy targeting indoleamine 2, 3-dioxygenase 1 (IDO) activates innate immunity and mitigates colorectal cancer growth. Cancer Gene Ther 2020, 27(3):235–245.

45. Blache CA, Manuel ER, Kaltcheva TI, Wong AN, Ellenhorn JD, Blazar BR, Diamond DJ: Systemic delivery of Salmonella typhimurium transformed with IDO shRNA enhances intratumoral vector colonization and suppresses tumor growth. Cancer Res 2012, 72(24):6447–6456.

46. Manuel ER, Chen J, D’Apuzzo M, Lampa MG, Kaltcheva TI, Thompson CB, Ludwig T, Chung V, Diamond DJ: <em>Salmonella</em>-Based Therapy Targeting Indoleamine 2,3-Dioxygenase Coupled with Enzymatic Depletion of Tumor Hyaluronan Induces Complete Regression of Aggressive Pancreatic Tumors. Cancer Immunology Research 2015, 3(9):1096–1107.

47. Ebelt ND, Zuniga E, Passi KB, Sobocinski LJ, Manuel ER: Hyaluronidase-Expressing Salmonella Effectively Targets Tumor-Associated Hyaluronic Acid in Pancreatic Ductal Adenocarcinoma. Mol Cancer Ther 2019:molcanther.0556.2019.

48. Lane MC, Alteri CJ, Smith SN, Mobley HLT: Expression of flagella is coincident with uropathogenic <em>Escherichia coli</em> ascension to the upper urinary tract. Proceedings of the National Academy of Sciences 2007, 104(42):16669–16674.

49. Manuel ER, Blache CA, Paquette R, Kaltcheva TI, Ishizaki H, Ellenhorn JD, Hensel M, Metelitsa L, Diamond DJ: Enhancement of cancer vaccine therapy by systemic delivery of a tumor-targeting Salmonella-based STAT3 shRNA suppresses the growth of established melanoma tumors. Cancer Res 2011, 71(12):4183–4191.

50. Phan T, Nguyen VH, D’Alincourt MS, Manuel ER, Kaltcheva T, Tsai W, Blazar BR, Diamond DJ, Melstrom LG: Salmonella-mediated therapy targeting indoleamine 2, 3-dioxygenase 1 (IDO) activates innate immunity and mitigates colorectal cancer growth. Cancer Gene Ther 2019.

51. Kim TS, Braciale TJ: Respiratory Dendritic Cell Subsets Differ in Their Capacity to Support the Induction of Virus-Specific Cytotoxic CD8+ T Cell Responses. PLoS One 2009, 4(1):e4204.

52. Jakubzick C, Gautier EL, Gibbings SL, Sojka DK, Schlitzer A, Johnson TE, Ivanov S, Duan Q, Bala S, Condon T et al: Minimal differentiation of classical monocytes as they survey steady-state tissues and transport antigen to lymph nodes. Immunity 2013, 39(3):599–610.

53. Hey YY, O’Neill HC: Murine spleen contains a diversity of myeloid and dendritic cells distinct in antigen presenting function. J Cell Mol Med 2012, 16(11):2611–2619.

54. Yu P, Lee Y, Liu W, Chin RK, Wang J, Wang Y, Schietinger A, Philip M, Schreiber H, Fu Y-X: Priming of naive T cells inside tumors leads to eradication of established tumors. Nat Immunol 2004, 5(2):141–149.

55. Kim H-D, Song G-W, Park S, Jung MK, Kim MH, Kang HJ, Yoo C, Yi K, Kim KH, Eo S et al: Association Between Expression Level of PD1 by Tumor-Infiltrating CD8+ T Cells and Features of Hepatocellular Carcinoma. Gastroenterology 2018, 155(6):1936-1950.e1917.

56. Ovcinnikovs V, Ross EM, Petersone L, Edner NM, Heuts F, Ntavli E, Kogimtzis A, Kennedy A, Wang CJ, Bennett CL et al: CTLA-4–mediated transendocytosis of costimulatory molecules primarily targets migratory dendritic cells. Science Immunology 2019, 4(35):eaaw0902.

57. Hobo W, Maas F, Adisty N, de Witte T, Schaap N, van der Voort R, Dolstra H: siRNA silencing of PD-L1 and PD-L2 on dendritic cells augments expansion and function of minor histocompatibility antigen–specific CD8+ T cells. Blood 2010, 116(22):4501–4511.

58. Kamada T, Togashi Y, Tay C, Ha D, Sasaki A, Nakamura Y, Sato E, Fukuoka S, Tada Y, Tanaka A et al: PD-1<sup>+</sup> regulatory T cells amplified by PD-1 blockade promote hyperprogression of cancer. Proceedings of the National Academy of Sciences 2019, 116(20):9999–10008.

59. Dodagatta-Marri E, Meyer DS, Reeves MQ, Paniagua R, To MD, Binnewies M, Broz ML, Mori H, Wu D, Adoumie M et al: α-PD-1 therapy elevates Treg/Th balance and increases tumor cell pSmad3 that are both targeted by α-TGFβ antibody to promote durable rejection and immunity in squamous cell carcinomas. Journal for ImmunoTherapy of Cancer 2019, 7(1):62.

60. Buchan SL, Fallatah M, Thirdborough SM, Taraban VY, Rogel A, Thomas LJ, Penfold CA, He L-Z, Curran MA, Keler T et al: PD-1 Blockade and CD27 Stimulation Activate Distinct Transcriptional Programs That Synergize for CD8<sup>+</sup> T-Cell–Driven Antitumor Immunity. Clin Cancer Res 2018, 24(10):2383–2394.

61. Yang S, Liu F, Wang QJ, Rosenberg SA, Morgan RA: The Shedding of CD62L (L-Selectin) Regulates the Acquisition of Lytic Activity in Human Tumor Reactive T Lymphocytes. PLoS One 2011, 6(7):e22560.

62. Hodge G, Barnawi J, Jurisevic C, Moffat D, Holmes M, Reynolds PN, Jersmann H, Hodge S: Lung cancer is associated with decreased expression of perforin, granzyme B and interferon (IFN)-γ by infiltrating lung tissue T cells, natural killer (NK)?T-like and NK cells. Clin Exp Immunol 2014, 178(1):79–85.

63. Ni L, Lu J: Interferon gamma in cancer immunotherapy. Cancer Med 2018, 7(9):4509–4516.

64. Krijgsman D, Hokland M, Kuppen PJK: The Role of Natural Killer T Cells in Cancer—A Phenotypical and Functional Approach. Front Immunol 2018, 9(367).

65. Takashima A, Yao Y: Neutrophil plasticity: acquisition of phenotype and functionality of antigen-presenting cell. J Leukocyte Biol 2015, 98(4):489–496.

66. Nelson CE, Mills LJ, McCurtain JL, Thompson EA, Seelig DM, Bhela S, Quarnstrom CF, Fife BT, Vezys V: Reprogramming responsiveness to checkpoint blockade in dysfunctional CD8 T cells. Proceedings of the National Academy of Sciences 2019, 116(7):2640–2645.

67. Pauken KE, Nelson CE, Martinov T, Spanier JA, Heffernan JR, Sahli NL, Quarnstrom CF, Osum KC, Schenkel JM, Jenkins MK et al: Cutting edge: identification of autoreactive CD4+ and CD8+ T cell subsets resistant to PD-1 pathway blockade. J Immunol 2015, 194(8):3551–3555.

68. Maeda Y, Nishikawa H, Sugiyama D, Ha D, Hamaguchi M, Saito T, Nishioka M, Wing JB, Adeegbe D, Katayama I et al: Detection of self-reactive CD8(+) T cells with an anergic phenotype in healthy individuals. Science 2014, 346(6216):1536–1540.

69. Philip M, Fairchild L, Sun L, Horste EL, Camara S, Shakiba M, Scott AC, Viale A, Lauer P, Merghoub T et al: Chromatin states define tumour-specific T cell dysfunction and reprogramming. Nature 2017, 545(7655):452–456.

70. Berzofsky JA, Terabe M: NKT Cells in Tumor Immunity: Opposing Subsets Define a New Immunoregulatory Axis. The Journal of Immunology 2008, 180(6):3627–3635.

71. Terabe M, Berzofsky JA: Tissue-Specific Roles of NKT Cells in Tumor Immunity. Front Immunol 2018, 9(1838).

72. Nair S, Dhodapkar MV: Natural Killer T Cells in Cancer Immunotherapy. Front Immunol 2017, 8(1178).

73. Ihara F, Sakurai D, Takami M, Kamata T, Kunii N, Yamasaki K, Iinuma T, Nakayama T, Motohashi S, Okamoto Y: Regulatory T cells induce CD4-NKT cell anergy and suppress NKT cell cytotoxic function. Cancer Immunol, Immunother 2019, 68(12):1935–1947.

74. Fallarino F, Grohmann U, You S, McGrath BC, Cavener DR, Vacca C, Orabona C, Bianchi R, Belladonna ML, Volpi C et al: The combined effects of tryptophan starvation and tryptophan catabolites down-regulate T cell receptor zeta-chain and induce a regulatory phenotype in naive T cells. J Immunol 2006, 176(11):6752–6761.

75. Maria NI, van Helden-Meeuwsen CG, Brkic Z, Paulissen SM, Steenwijk EC, Dalm VA, van Daele PL, Martin van Hagen P, Kroese FG, van Roon JA et al: Association of Increased Treg Cell Levels With Elevated Indoleamine 2,3-Dioxygenase Activity and an Imbalanced Kynurenine Pathway in Interferon-Positive Primary Sjogren’s Syndrome. Arthritis Rheumatol 2016, 68(7):1688–1699.

76. Quintana FJ, Murugaiyan G, Farez MF, Mitsdoerffer M, Tukpah A-M, Burns EJ, Weiner HL: An endogenous aryl hydrocarbon receptor ligand acts on dendritic cells and T cells to suppress experimental autoimmune encephalomyelitis. Proceedings of the National Academy of Sciences 2010, 107(48):20768–20773.

77. Thatcher TH, Williams MA, Pollock SJ, McCarthy CE, Lacy SH, Phipps RP, Sime PJ: Endogenous ligands of the aryl hydrocarbon receptor regulate lung dendritic cell function. Immunology 2016, 147(1):41–54.

