## Supplementary Figures for "Restructuring of the Immune Contexture Improves Checkpoint Blockade Efficacy in Murine Lung Cancer"

Supplemental Material

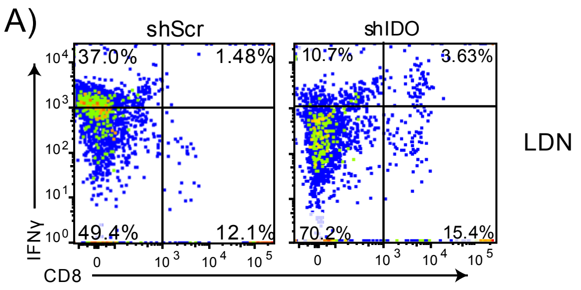


**Supplemental Figure 1. Phenotypes of SIINFEKL-specific CD8+ splenocytes after incubation with neutrophils induced by shScr or shID-ST. (A)** Flow cytometry dot plots of activated CD8 T cells (CD8 and IFNγ double positive) out of total OTI splenocytes after 24 hours of *in vitro* co-incubation with fixed, SIINFEKL-loaded LDN from either shScr or shIDO-ST treated mice.

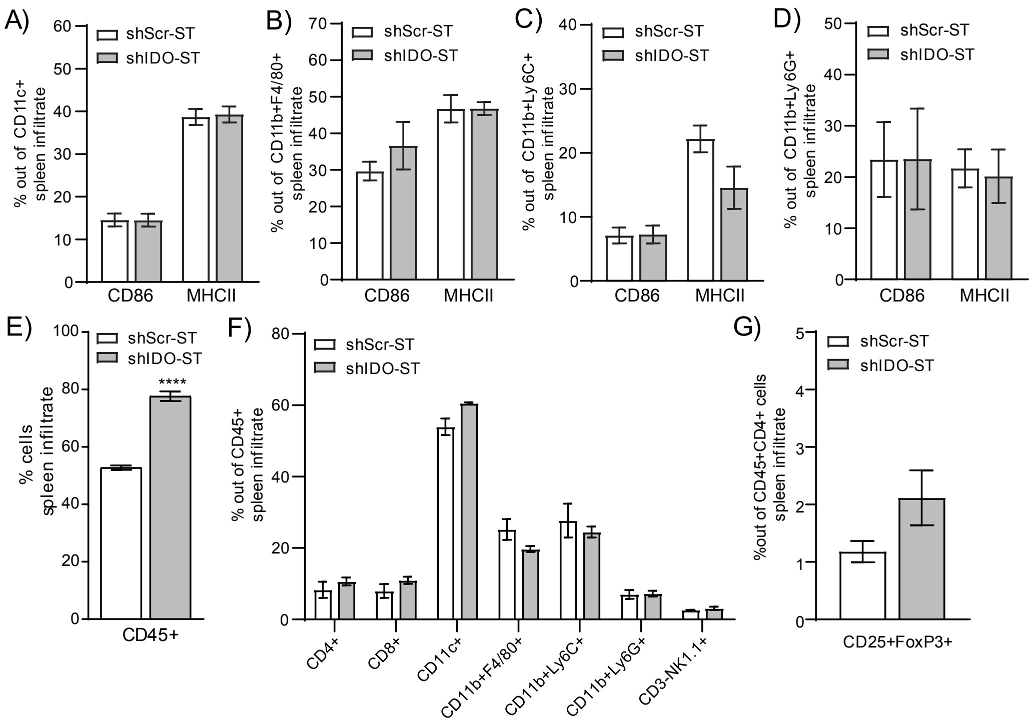


**Supplemental Figure 2. Phenotypes and frequencies of splenocytes in tumor-bearing mice after shIDO-ST treatment.** Six days after implantation of LLC1 cells into mice, mice were treated with three consecutive, daily doses of $1x10^6$ cfu shScr or shIDO-ST. Spleens were processed 48 hours after the third treatment. All bar graphs represent quantifications of flow cytometry. Percentages of CD86 or MHCII positive cells by immune cell type were quantified **(A)** out of CD11c+ cells, **(B)** out of CD11b+F4/80+ cells, **(C)** out of CD11b+Ly6C+ cells, and **(D)** out of CD11b+Ly6G+ cells. **(E)** Total CD45+ cells and **(F)** individual cell types out of total CD45+ splenocytes were quantified. **(G)** The percentage of splenic Tregs (CD25+Foxp3+) was quantified out of CD45+CD4+ cells. *p<0.05, **p<0.01, ***p<0.001, ****p<0.0001.

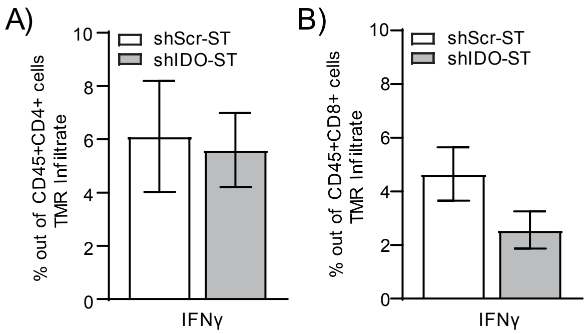


**Supplemental Figure 3. Activation of intratumoral T cells after shIDO-ST treatment.** Six days after implantation of LLC1 cells into mice, mice were treated with three consecutive, daily doses of $1x10^6$ cfu shScr or shIDO-ST. Tumors were processed 48 hours after the third treatment. All bar graphs represent quantifications of flow cytometry. Percentages of IFNγ positive cells by immune cell type were quantified **(A)** out of $CD4^+$ cells and **(B)** out of $CD8^+$ cells.
